# FracFixR: A compositional statistical framework for absolute proportion estimation between fractions in RNA sequencing data

**DOI:** 10.1101/2025.07.29.667459

**Authors:** Alice Cleynen, Agin Ravindran, Nikolay Shirokikh

## Abstract

**Motivation:** RNA fractionation followed by sequencing is widely used to study RNA localization, translation, and subcellular compartmentalization. Interpreting fractionated RNA-seq data poses a fundamental compositional challenge: library preparation and sequencing depth obscure the original proportions of RNA fractions, which can bias comparisons – particularly when biological changes shift RNA distribution across fractions. This bias compromises comparisons of fraction-specific RNA profiles and limits the utility of standard differential expression methods. Existing approaches using transcript frequency ratios or standard normalization fail to account for the compositional nature of fractionated samples and cannot estimate the unrecoverable “lost” fraction.

**Results:** We developed FracFixR, a statistical framework that reconstructs original fraction proportions by modeling the compositional relationship between whole and fractionated RNA samples. Using non-negative linear regression on carefully selected transcripts, FracFixR estimates global fraction weights, corrects individual transcript frequencies, and quantifies unrecoverable material. The framework includes methods for differential proportion testing between conditions using binomial GLM, logit, or beta-binomial models. We rigorously validated FracFixR using synthetic data with known ground truth and real polysome profiling data from multiple cell lines, demonstrating accurate reconstruction of fraction weights (Pearson correlation > 0.85) and enabling detection of differentially translated transcripts between cancer subtypes.

**Availability and implementation:** FracFixR is implemented as an R package freely available on GitHub at https://github.com/Arnaroo/FracFixR.

## 1 Introduction

Understanding the functional state of a cell often begins with a snapshot of its transcriptome. RNA sequencing (RNA-seq) has become a powerful and widely adopted tool to capture such snapshots, enabling researchers to quantify and compare transcript abundances across different biological conditions [8, 29]. To gain a more refined understanding of gene expression control, many experimental protocols now include fractionation steps before sequencing. For example, separating cytoplasmic and nuclear RNA can reveal regulatory mechanisms that govern nuclear export and stability of RNA [7, 28], while polysome profiling can isolate ribosome-associated transcripts to focus on translated mRNA and identify their ribosomal load [9, 6].

However, interpreting fractionated sample RNA sequencing data presents a unique compositional challenge not adequately addressed by standard differential expression analysis methods [24]. This occurs because library preparation steps – possibly including amplification of the initial material – can lead to a loss of information about the original fraction proportions (Figure 1a), especially when the entire sample is not captured due to sequencing depth limitations and inability to collect all fractions of the whole. These effects bias and cripple any comparisons of fraction-specific RNA profiles, which are often the major goal of the experiment. The bias may be further inflated by global biological shifts in RNA distribution across the fractions when comparing different conditions (such as stress or drug response), where much RNA can end up in non-recoverable “lost” fraction [12, 6].

**Figure 1.**
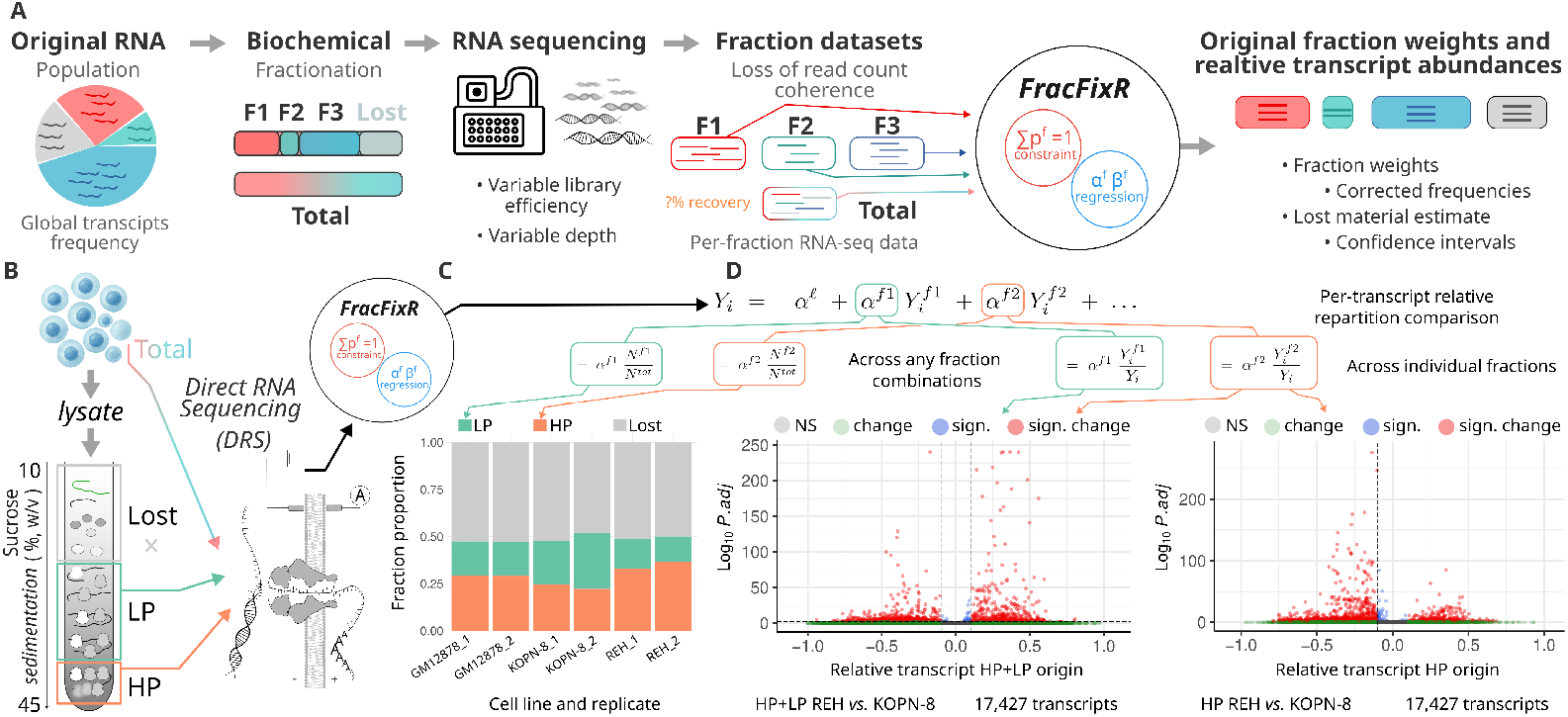
FracFixR: a convenient tool to enable cross-fraction comparisons of relative transcript abundances and recover the original fraction weights in RNA-seq data. (**A**)FracFixR addresses the different sequencing depth, library efficiency and non-sequenced “lost” material problems of a typical RNA-sequencing experiment of fractionated cell material. (**B**) Polysome profiling with separation into heavy polysomes (HP) and light polysomes (LP) as a generic RNA fractionation experiment. (**C**) FracFixR reconstructs polysome profiling fraction weights and calculates the unobserved “lost” fraction in an experiment with differentlyrelapsing B-ALL cell lines REH and KOPN-8, and a non-malignant B-cell control (GM12878). (**D**) FracFixR reconstructs individual transcript relative abundance profiles across the fractions and detects many substantial and significant translationally-controlled mRNAs between REH and KOPN8.

Of the common techniques, polysome to monosome ratios (P:M [10]) fails to distinguish the free from translating ribosomes, while transcript frequency-based ratios (such as the Polysome Sedimentation Factor, PSF) can be insufficient because they assume constant global fraction proportions, which is violated when the overall distribution into fractions changes (such as the cell translated overall more or less RNA [12]). Differential transcript abundance (DTA) analysis applied directly across fractions (such as DESeq2 [18]) is inappropriate because it ignores that fractions are parts of a whole. Standard normalization is also inadequate because it assumes proportional scaling, which is violated when recovery patterns differ [4, 23].

The complexity of the transcriptome, with its extensive non-coding RNA components and intricate regulatory networks [19, 20, 21], further compounds these analytical challenges. RNA localization and compartmentalization are increasingly recognized as fundamental aspects of gene regulation [22, 11, 5], making accurate quantification of RNA distribution across cellular fractions essential for understanding cellular functions.

Here we introduce `FracFixR`, a general-purpose statistical framework for analyzing RNA-seq data obtained from any fractionation protocol. FracFixR reconstructs the proportions of fractions lost by analyzing data from all fractions in an integrated model, estimates global and transcript-level shifts from available data, and provides best possible approximation given information constraints (Figure 1a). `FracFixR` takes matrix of the mapped read counts as the input and performs compositional fraction normalization. `FracFixR` outputs: (a) restored weights of original fractions, (b) frequencies of each transcript, and (c) weight of unrecoverable material.

`FracFixR` can be broadly applied to any RNA fractionation experiment – from subcellular localization studies [27] to ribosome profiling [13, 17, 25, 1] to RNA-protein complex isolation [2, 26] – enabling researchers to confidently detect biological changes in RNA distribution patterns across diverse experimental systems.

### 2 FracFixR implementation

#### 2.1 Development of the FracFixR statistical model

The statistical model is based on the compositional relationship between the whole and the fractions of the RNAs in the cell.

##### Notations

In what follows, we denote by *j* ∈ {1, *n*} as the original whole, from which a set of fractions are derived and sequenced. For simplicity, we will denote ℱ the set of all sequenced fractions and its unobserved or “lost’ complement, and we will denote *f* an element of ℱ (quantities not indexed by any fraction will represent quantities in the total population). Finally, *i* ∈ {1 : *G*} denotes a transcript (or a gene), and 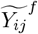 denotes the true (but unobserved) number of transcripts *i* in fraction *f* of sample *j*, while *Y*_*ij*_^*f*^ will denote the number of observed counts.

##### Global fractions

A common assumption is that *Y*_*ij*_^*f*^ is a direct representation of 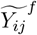 up to the amplification and sampling effects, and thus we assume 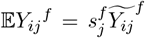. Furthermore, for any individual *j* and transcript *i*, we introduce 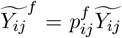 such that 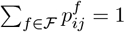.

The relationships between fractions and the total population lead to

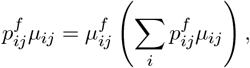

where 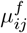 is the proportion of transcript *i* in fraction *f* of individual *j*. We estimate the coefficients 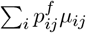, representing the global proportion of fraction *f* in individual *j*, using non-negative least squares (NNLS) regression. The model relates total and fraction-specific counts via:

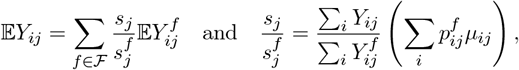

This leads to the estimate:

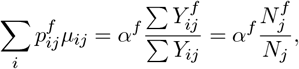

where *α*^*f*^ is the NNLS coefficient for fraction *f*, wand the unobserved fraction is captured by the intercept.

##### Individual transcripts

To estimate individual transcript proportions 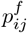 in each fraction, we then use the contribution of fraction *f* in the total count *Y*_*ij*_ *via* equation

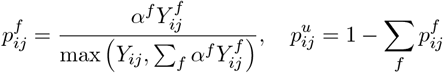

where the max function ensures that all proportions are positive.

To compare true relative abundances of individual transcripts across a given fraction (or set of fractions) between two conditions, we introduce a binomial generalized linear model:

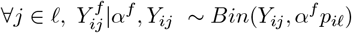

and test 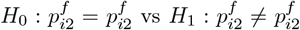. P-values are adjusted for multiple testing with the Benjamini-Hochberg procedure [3].

We also provide implementations of a beta-binomial Wald test and a logit-based test as faster alternatives, though they may offer reduced statistical power.

### 2.2 FracFixR: a convenient and user-friendly R package

`FracFixR` is implemented as an R package freely available on GitHub. Its standard workflow takes as input a matrix of raw transcript or gene counts from the total population, along with one or more sequenced fractions. It also requires an annotation data frame specifying sample names, experimental conditions, replicates, and fraction types. The package returns an object containing the original data, a matrix of fraction-corrected counts, estimated fraction abundances, normalization coefficients, and diagnostic regression plots.

`FracFixR` can be installed from GitHub using R command install.packages(“https://github.com/Arnaroo/FracFixR/releases/download/v1.0.0/FracFixR_1.0.0.tar.gz”, repos= NULL, type = “source”). It has modest dependencies, including the `nnls` package to perform the non-negative linear model, and requires standard data manipulation packages (`tidyr, dplyr` and `matrixStats`), plotting packages (`ggplot2, RColorBrewer, EnhancedVolcano`). `FracFixR` makes use of the `future.apply` package for parallel running of the glm test.

`FracFixR` is ran using two commands:

- `Norma<-FracFixR(CountMatrix,Annotation)` runs the NNLS regession and estimates global fractions and individual contributions. The global fraction estimation can be visualized using the command PlotFractions(Norma).
- Results<-DiffPropTest(Norma,Conditions=c(“Condition1”,”Condition2”),Types = “Fraction1”, Test = “GLM”) performs the comparisons between conditions. The resulting volcano plot can be visualized using the command PlotComparison(Results).

Detailed instructions and readme are available in the FracFixR GitHub repository.

## 3 FracFixR performance

In any fractionated RNA-seq experiment, the original ground truth is inherently lost. To rigorously evaluate the performance of FracFixR, we used specialized datasets with either synthetic ground truth or mixtures of real data with known fraction proportions (see Methods for details). We then applied FracFixR to real data obtained from long-read direct RNA sequencing (DRS) of the GM12878 Epstein-Barr virus-immortalized normal B-cell line, as well as the B-cell acute lymphoblastic leukemia (B-ALL) REH and KOPN-8 cell lines, to compare their translation efficiency. For each cell line, total RNA was extracted under denaturing lysis conditions, while fractionated RNA was obtained from light and heavy polysomal fractions following sucrose gradient sedimentation (see Methods).

### 3.1 FracFixR validation and testing

To construct a ground truth and evaluate modeling accuracy across diverse scenarios, we began with a real count distribution from a DRS experiment containing 1.8 million total reads. We adjusted the total read count to the desired level by subsampling or scaling, then partitioned the data into two observable fractions and a “lost” fraction. The two fractions were allowed to differ in their global representation within the total RNA pool, and transcript abundance profiles within each fraction were varied to reflect distinct biological behaviors. These synthetic datasets were further subsampled to simulate varying recovery rates in both total and fractionated libraries, mimicking different levels of RNA loss and sequencing depth (Figure 2). This design captures a range of realistic and challenging conditions, including imbalanced fraction proportions, substantial lost fractions, and uneven recovery or sequencing depth across fractions.

**Figure 2.**
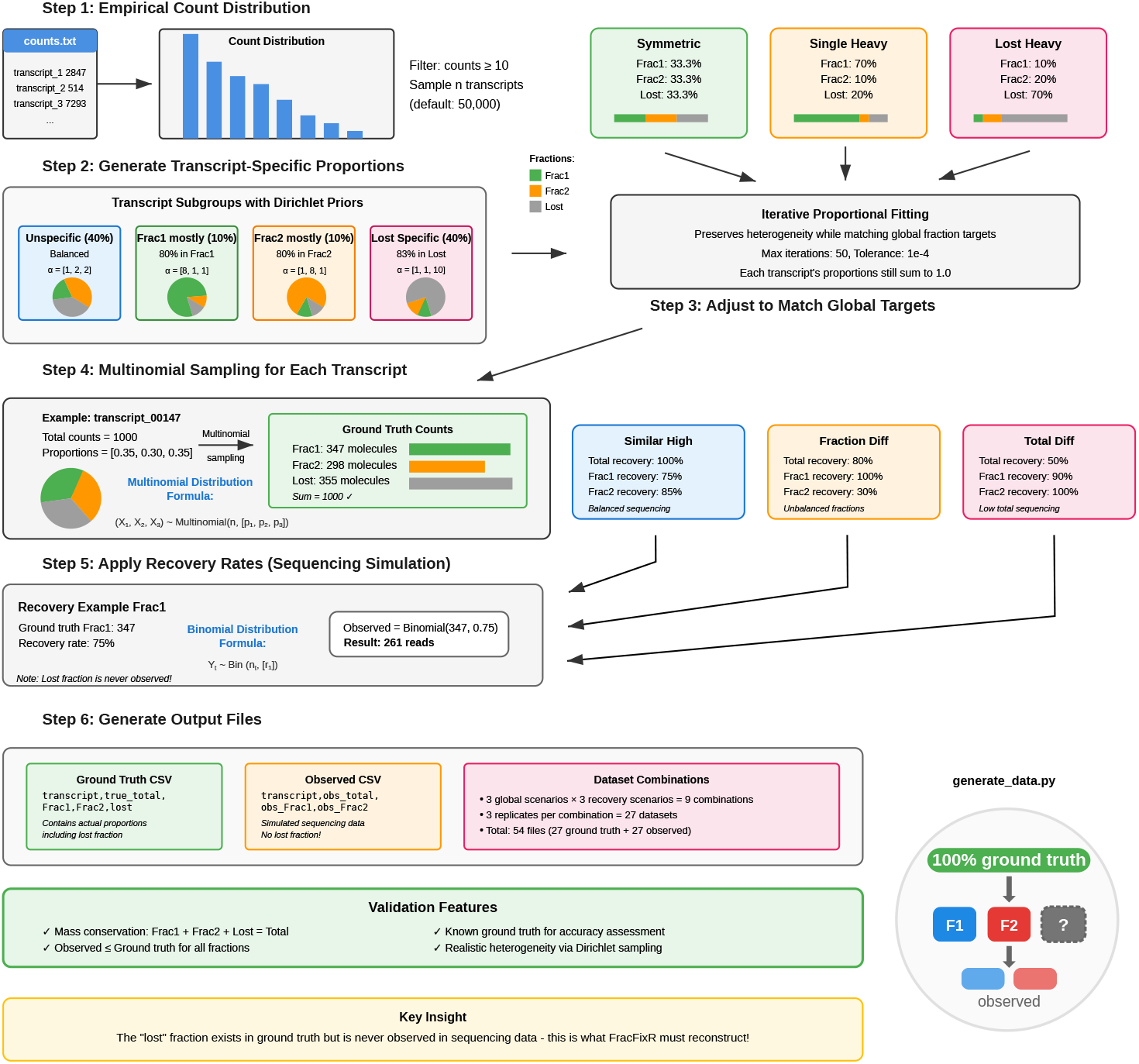
Synthetic data generation workflow for FracFixR validation. Step 1 begins with empirical count distribution from real RNA-seq data, filtering transcripts with counts greater than or equal to 10 and sampling n transcripts (default 50,000). Step 2 assigns transcripts to four subgroups with distinct fractionation behaviors using different Dirichlet prior distributions. Step 3 applies iterative proportional fitting to adjust individual transcript proportions while maintaining heterogeneity and matching global fraction targets. Step 4 implements three global scenarios representing different biological conditions. Step 5 uses multinomial sampling to distribute each transcript’s total counts across fractions according to its specific proportions, ensuring mass conservation. Step 6 simulates sequencing by applying recovery rates through binomial sampling, testing three sequencing-depth (recovery) scenarios. Step 7 generates output files: ground truth CSVs containing all fractions including lost material, and observed CSVs containing only sequenceable fractions, mimicking real experimental data where lost material cannot be observed.

FracFixR consistently and accurately recovered global fraction weights, including the “lost” fraction, across all simulated scenarios, with Pearson correlations near 1, RMSE *<* 0.17, and dispersion *<* 0.15 – even under challenging conditions with asymmetry or data loss (Figure 3). Transcript-level abundances also closely matched ground truth across all fractions, confirming the statistical validity of the framework. Interestingly, the precision in the estimation of the transcript-level abundance is directly related to the recovery rate (sequencing depth) of the fraction, and is almost insensitive to the actual true global fraction proportions, suggesting that deep sequencing should allow precise recovery of the abundances no matter what the exact fraction importance.

**Figure 3.**
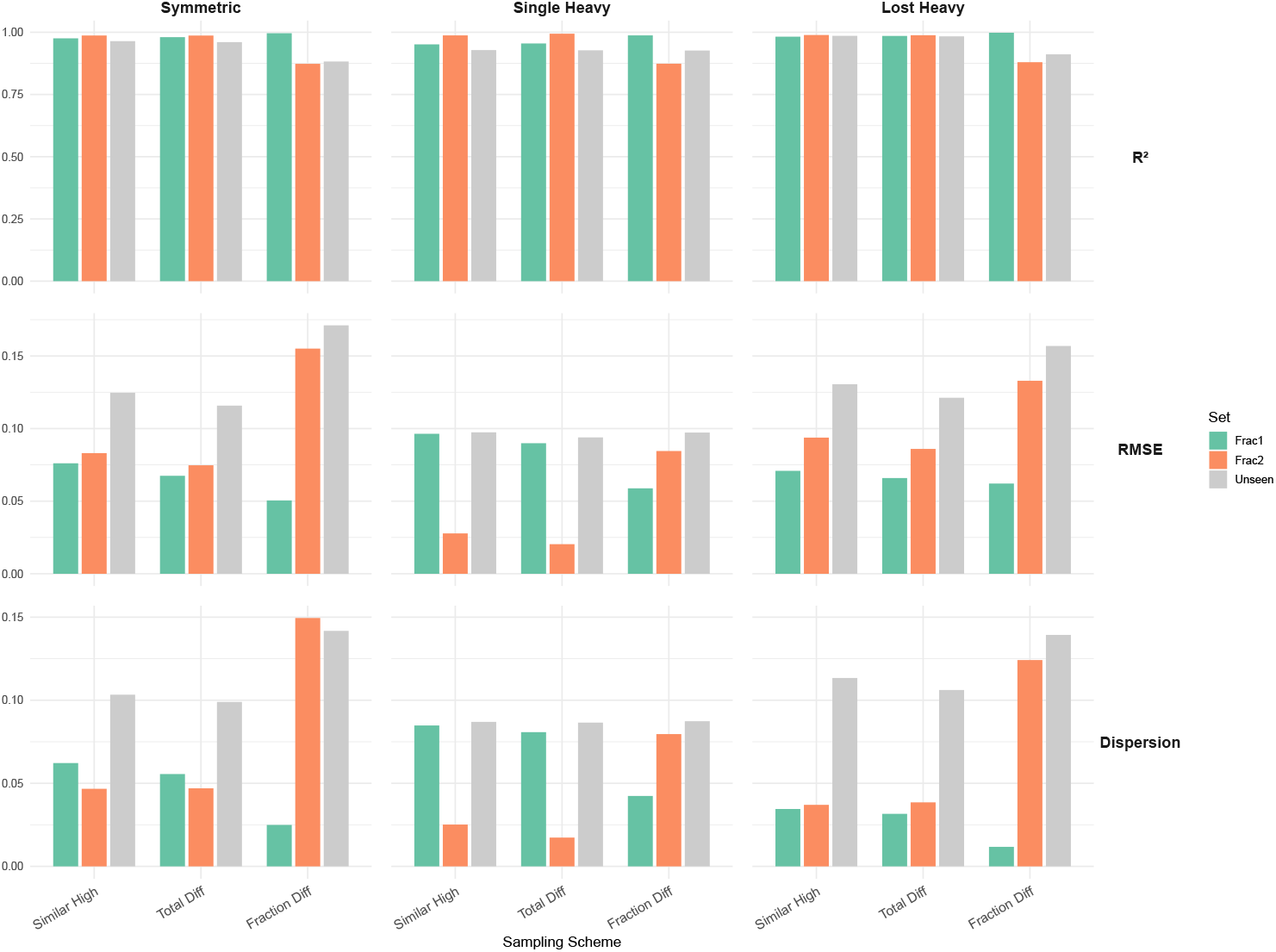
Comprehensive performance metrics of FracFixR across all synthetic validation scenarios. Box plots show *R*^2^ (coefficient of determination), RMSE (root mean square error), and dispersion values for each fraction (Frac1, Frac2, Unseen) across nine experimental conditions combining three proportion scenarios (Symmetric, Single Heavy, Lost Heavy) with three recovery schemes (Similar High, Total Diff, Fraction Diff). High *R*^2^ values (*>* 0.85) and low RMSE (*<* 0.15) across all conditions demonstrate robust fraction weight reconstruction even under challenging recovery patterns.

To understand how the performance expectations could translate to data derived from an actual sequencing experiment, we additionally performed three controlled simulations from actual fastq reads, mixing RNA-seq of the total cell lysate of REH, KOPN-8 and GM cells, as well as polysomal fractions of REH and KOPN-8 with GM total cell lysate as the “unseen” fraction. The mixing ratios were variable but known. Upon mapping, only the subsets of total mix, REH and KOPN-8 were presented to FracFixR, with GM reads playing the role of a lost “unobserved” fraction. Once again, FracFixR was able to accurately reconstruct the original fraction proportions (weights), as well as provide an estimate of the amount of the “unknown” lost fraction (Figure 4), proving it can handle the biases and variability associated with real data.

**Figure 4.**
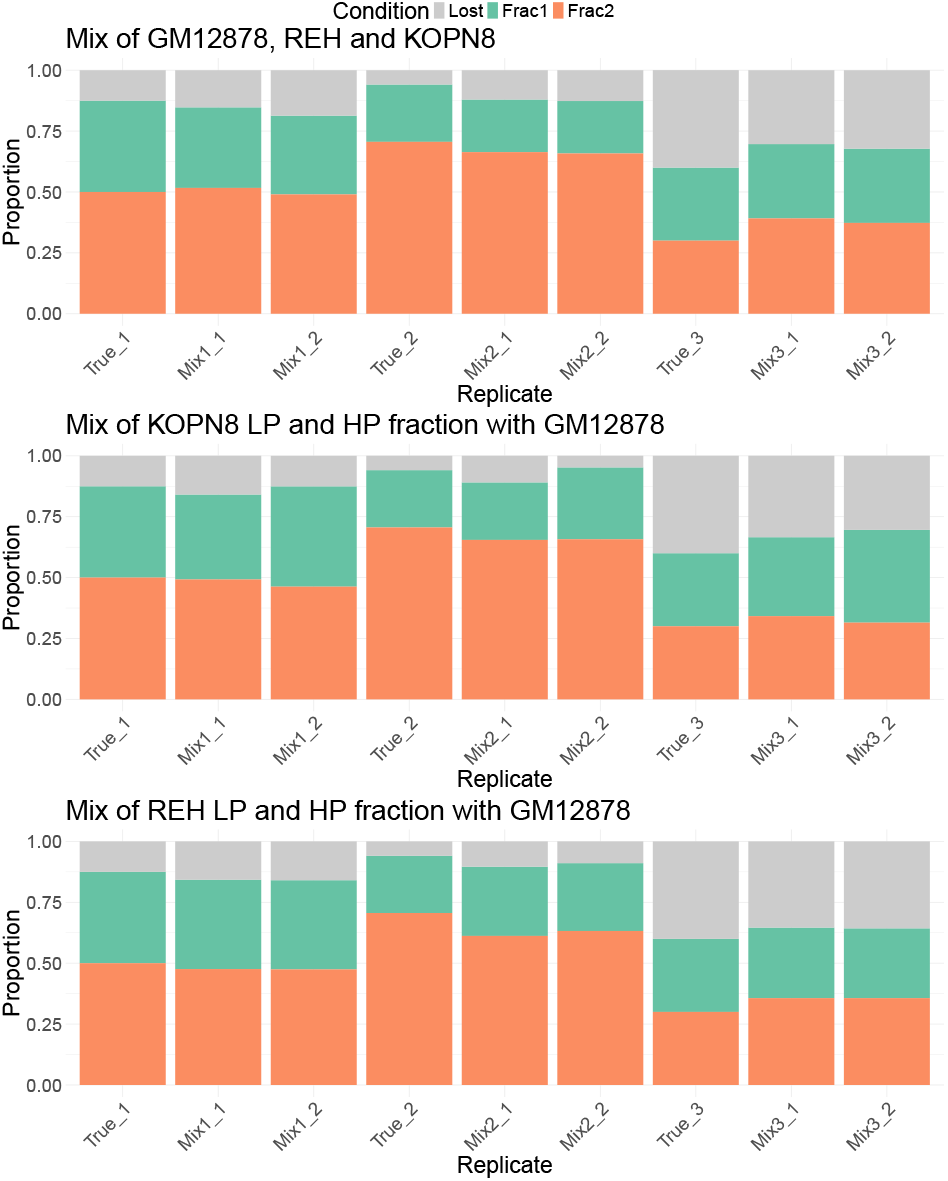
FracFixR reconstruction of fraction weights in controlled mixing experiments using real RNA-seq data. Stacked bar plots show the estimated proportions of lost (grey), Frac1 (orange), and Frac2 (green) fractions for three mixing scenarios: (top) equal mix of GM12878, REH, and KOPN-8 total cell lysates; (middle) mix of KOPN-8 polysomal fractions with GM12878 total lysate; (bottom) mix of REH polysomal fractions with GM12878 total lysate. True_N represents known mixing proportions, while Mix_N represents FracFixR estimates from observed data only in two different replicates, demonstrating accurate recovery of hidden GM12878 fraction as “lost” material.

### 3.2 FracFixR substantially corrects relative transcript abundances in fractionated RNA data

We then applied FracFixR to a real polysome profiling dataset derived from three cell lines: GM (non-cancerous), REH, and KOPN-8 (both B-ALL subtypes with distinct relapse potential). For each line, we obtained RNA from total lysates as well as from cytoplasmic material fractionated into light (1–3 ribosomes) and heavy (≥ 4 ribosomes) polysomes (Figure 1b). As is typical for such experiments, a considerable proportion of RNA was not captured in the sequenced data, due to material loss during lysis and exclusion of non-polysomal fractions.

`FracFixR` reconstructed the true composition of each sample, including the unsequenced “lost” fraction, and revealed distinct patterns of translation engagement across cell lines. As shown before, a vast majority of transcripts were not associated with ribosomes [14, 10] GM and REH cells showed greater representation of transcripts in heavy polysomes compared to KOPN-8, indicating a higher overall level of translational activity (Figure 1c).

Using `FracFixR`, we further were able to compare the per-transcript differences between the REH and KOPN-8 cell lines, revealing many transcripts with cell-line specific control. In the combined polysome association analysis enabled by FracFixR, 406 transcripts were more associated with ribosomes in KOPN-8, and 1,335 transcripts were more associated with ribosomes in REH (Figure 1D, Figure 5). Gene ontology of these hits revealed distinct enriched control pathways. Transcripts showing increased polysome association in REH cells are enriched for metabolic pathways, particularly biosynthesis of amino acids (fold enrichment 6), carbon metabolism, and glycolysis/gluconeogenesis. Notably, the spliceosome pathway shows significant enrichment, suggesting coordinated translational upregulation of RNA processing machinery. Central carbon metabolism in cancer and nucleocytoplasmic transport pathways are also enriched, indicating enhanced metabolic activity and cellular transport processes. KOPN-8 cells show striking enrichment for ribosome components (fold enrichment 30), indicating substantial translational control of the protein synthesis machinery itself. This suggests a feed-forward mechanism where increased translation capacity is achieved through preferential translation of ribosomal proteins. Additional enrichment in oxidative phosphorylation and metabolic pathways indicates distinct metabolic programming compared to REH cells. The differential enrichment patterns reveal cell line-specific translational control mechanisms: REH cells prioritize amino acid biosynthesis and RNA processing, while KOPN-8 cells enhance their translational capacity through ribosomal protein synthesis. Gene lists passing the significance and effect size cut-offs from the differential comparison engine of FracFixR (GLM) were further analyzed by ShinyGO 0.82 in the KEGG database.

**Figure 5.**
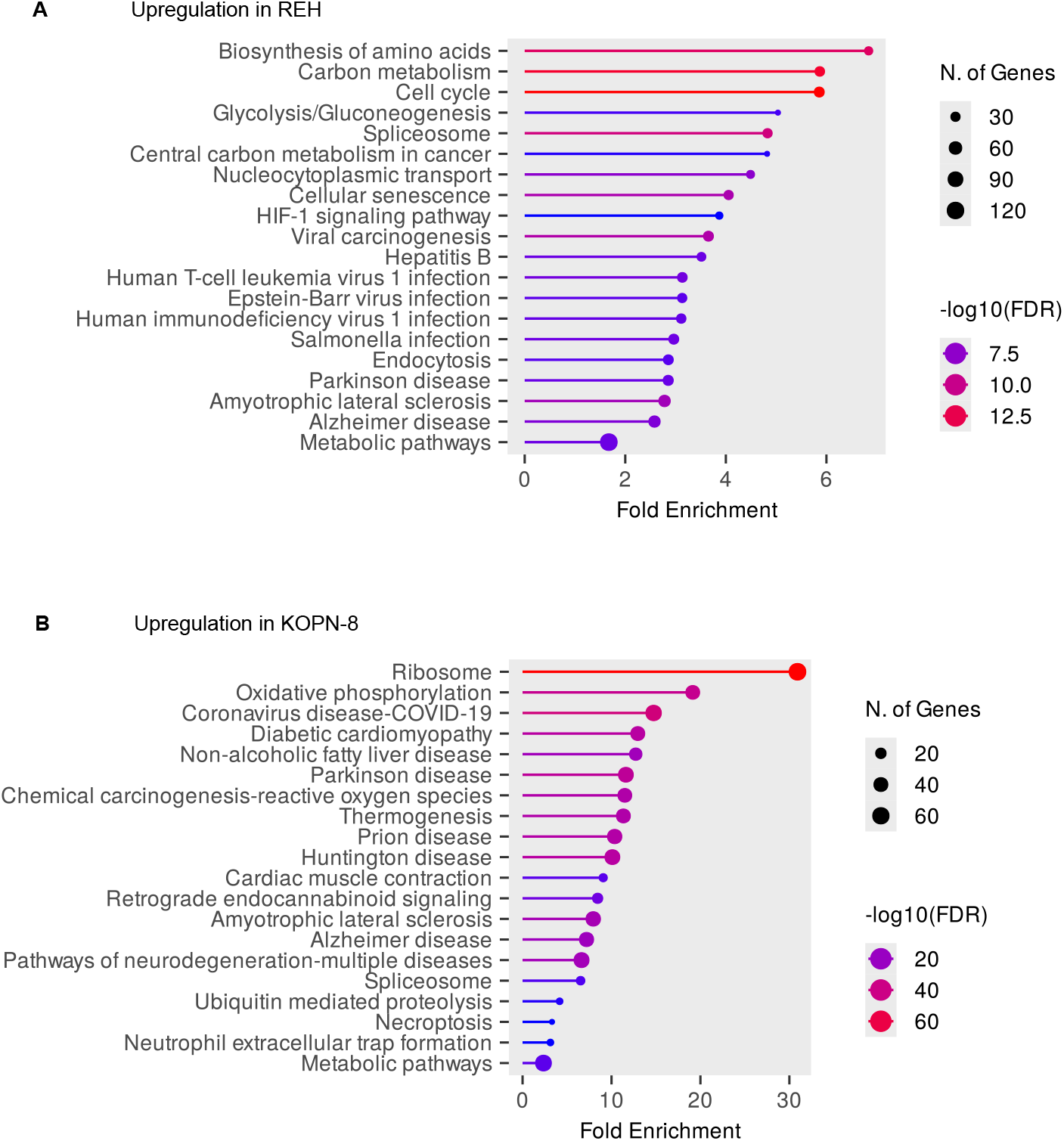
Pathway enrichment analysis reveals distinct translational programs between REH and KOPN-8 B-ALL cell lines. Dot size represents the number of genes in each pathway, color intensity indicates -log10(FDR), with darker colors representing more significant enrichment.

## 4 Materials and Methods for model cell-lines

### 4.1 Cell lines and growth conditions

The B-ALL fusion-driven cell lines, REH and KOPN-8 cell lines were purchased from Leibniz Institute DSMZ, Germany. The non-B-ALL control cell line, GM12878 cell line was purchased from Coriell Institute for Medical Research, USA. All three cell lines were cultured under controlled conditions at 37°C and 5% CO2. The cells were cultured in RPMI-1640 media (Sigma-Aldrich), supplemented with sodium bicarbonate and devoid of L-glutamine. Additionally, fetal bovine serum (FBS) was added to the RPMI-1640 media to a final concentration of 10%. Cells were split in a 1:4 split and maintained at a cell density of 2 million at ∼90 % confluency.

### 4.2 Total (whole cell) RNA extraction

For whole cell transcriptome analysis, each cell line was cultured in RPMI-1640 as described above. The cell pellets were collected when cells attained a minimum cell density of 2 × 10^6^ cells per ml of 50 ml media at a cell viability of at least 95 %. This ensured optimal cell viability and yield for all experimental procedures including downstream direct RNA sequencing and inhibitor based-polysome profiling. For the pellet collection, cells were pooled and spun down at 500 × g for 5 minutes. The supernatant was carefully discarded using a serological pipette, and the resulting cell pellet was washed with 1 mL of ice-cold phosphate-buffered saline (PBS). Following the PBS wash, cells were centrifuged again at 4°C for 5 minutes at 500 × g to ensure thorough pelleting, and the supernatant discarded. The PBS wash step was repeated after which the cell pellet was immediately processed further or stored at -80°C.

Total RNA was then isolated using the PureLink RNA Mini Kit (Thermo Fisher Scientific) following the manufacturer’s protocol. Briefly, cells were homogenised and lysed in lysis buffer, and RNA was selectively bound to the provided RNA-binding columns. Contaminants were removed through a series of wash steps, and purified RNA was eluted in RNase-free water. The quality and quantity of isolated RNA were assessed using a Nanodrop spectrophotometer (Thermo Fisher Scientific). This method ensured the extraction of high-quality RNA suitable for downstream applications such as direct RNA sequencing.

### 4.3 Translatome capture using polysome profiling

#### Translational cocktail inhibitor treatment and cell collection

Translational inhibitor cocktail, comprising of tigecycline, anisomycin and emetine, was administered to cells to preserve the translatome containing polysome-bound mRNA fractions. These prevalently utilised translational inhibitors bind to the elongating ribosome at distinct stages of the elongation cycle of protein synthesis. Emetine binds to the eukaryotic SSU (40S ribosomal subunit), while tigecycline binds to SSUs broadly, including the prokaryotic variant, and is also known to stall elongation, even within mitochondria. Anisomycin, on the other hand, binds to LSU (60S ribosomal subunit). This combined action of inhibitors was used to enhance elongation stalling efficiency compared to single drug interventions. The inclusion of this inhibitor cocktail stabilises translational and polysomal complexes during gradient fraction separation in polysome profiling, ensuring sufficient polysomal yield and a more authentic profile for the downstream direct RNA sequencing.

The antibiotic cocktail was prepared with Tigecycline (100 µL) added to a final concentration of 100 µg/mL, Emetine (10 µL) added to a final concentration of 20 µg/mL, and Anisomycin (133 µL) dissolved to a final concentration of 1 mM for every 20 mL of cells suspended in RPMI-1640 media. Following sequential addition of the premix to the wells and gentle pipetting, the cells were incubated until the 3-minute mark. Subsequently, the cells were pooled and centrifuged at 500 × g for 5 minutes. The resulting pellet was washed in 1 mL of 1× PBS and centrifuged again at 500 × g for 5 minutes at 4°C. The cell pellet was aspirated and stored at -80°C for subsequent cell lysis and gradient fractionation.

#### Cell Lysis and extraction

Lysis buffer was prepared by combining HEPES-KOH (pH 7.6) to a final concentration of 25 mM, KCl to a final concentration of 50 mM, DTT to 4 mM, MgCl_2_ to 5.1 mM, EDTA to 0.1 mM, Igepal CA-630 to 0.5%, glycerol to 5% (v/v), and emetine at 40 *µ*g/mL. All components were added and combined at stored at –80 °C, while DTT and EDTA were added only upon immediate use of the lysis buffer. The cell pellet, from overnight freezing at -80°C, was defrosted in ice and promptly treated with 1:2.5 ratio of pellet to lysis buffer supplemented with 1 *µ*L of 40 U/*µ*L RNAsin Plus (Promega). Subsequently, the cell suspension was vigorously pipette-mixed through a 200 *µ*L tip against a tube wall to avoid the introduction of air and foam formation. Further homogenisation was achieved by passing the suspension through 27.5G and then subsequently 31G syringes for up to 8-10 times each. Following this, the fully homogenised lysate was then centrifuged at 12,000 × g for 10 minutes at 4°C. The top layer containing the clarified lysate was collected avoiding cell debris material in the pellet, and stored at -80°C. Quality assessment of the lysate was performed by measuring optical density (OD) against that of the lysis buffer using a Nanodrop spectrophotometer.

#### Preparation of sucrose gradients

Light Sucrose (LS) Gradient Buffer was prepared by combining 25 mM HEPES-KOH (pH 7.6, 25°C), KCl to a final concentration of 100 mM, MgCl_2_ to 15 mM, EDTA to 0.1 mM, and a final concentration of 15% (w/v) sucrose. Heavy Sucrose (HS) gradient buffer was formulated with the same components as the LS Gradient Buffer, but sucrose concentration was elevated to 45% (w/v). For both LS and HS gradient buffers, emetine was added to a final concentration of 10 *µ*g/mL.

Sucrose buffer premixes were created by sequentially combining varying proportions of sucrose buffers and freshly added DTT to a final concentration of 10 mM. Each premix was stored on ice to preserve its reducing properties. Subsequently, the premixes were cast into Beckman-Coulter thin wall polypropylene tubes using an eppendorf dispenser preset to 2.5 ml. The casting process involved sequentially dispensing 2.5 ml of each premix into the tubes, starting from the highest sucrose concentration (45 %) sequentially to the lowest (15 %). The solution was layered into four distinct sections, each containing sucrose concentrations of 15 %, 25 %, 35 %, and 45 %, with each layer comprising 2.5 ml. After casting each layer, the tubes were promptly placed on dry ice for incremental freezing. This controlled freezing process ensured the formation of stable sucrose gradients within the tubes upon melting and reaching diffusional equilibration. The gradients were promptly stored at -80 °C. Before use, the tubes were transferred to a cold room to allow gradual thawing overnight prior to ultracentrifugation and gradient fractionation.

#### Gradient fractionation

Cell lysates were obtained after freezing at –80 °C freezer and were thawed on ice. Prior to the ultracentrifugation spin, each sucrose gradient centrifuge tube mass was measured and adjusted to ensure uniformity. Subsequently, a predetermined volume of cell lysate equivalent to an OD600-concentration of 50 OD units, was gently added to the top of each gradient, ensuring the pipette tip was submerged just beneath the surface of the 15 % sucrose layer and lysate was dispensed slowly minimising bubble formation. The prepared tubes were then placed within a chilled ultracentrifugation rotor and centrifuged for 2 hours at 45,000 rpm at 4 °C.

Upon completion, the tubes were carefully removed and placed in the cold room. Following this, 60% sucrose was run through the gradient fractionator, ensuring the absence of any air bubbles within the tubes. To establish baseline readings, the instrument was calibrated using an empty gradient without any cell lysate preloaded as a blank to ensure a stable baseline of the gradient is reached. This procedure was iteratively repeated for all subsequent sample gradients containing the cell lysate after ultracentrifugation. Finally, fractions containing light polysome (monosome, disome and trisome) sample tubes, and heavy polysome sample tubes (tetrasome, pentasome and onwards) were collected and preserved on dry ice, while the remaining portions were discarded.

#### Total RNA purification from polysome-bound fractions

Following the fractionation of polysomes, the fractions were pooled and combined into light and heavy polysomal fractions. To do so equal proportions of monosomes, disomes, and trisomes were combined and designated as the light polysome fraction, while resolved fractions of equal proportion from tetrasomes and higher order polysomes were pooled and labeled as the heavy polysome fraction. For RNA purification, EDTA was added to a final concentration of 50 mM and SDS to a final concentration of 2 % were added to each tube, followed by incubation at 55 °C for 5 minutes until the solution clarified. Subsequently, SPRI beads were resuspended, vortexed, and added to the sample at a 1:1 ratio of bead suspension to sample. After thorough mixing, the solution was incubated at room temperature for 5 minutes in a magnetic rack. The supernatant was discarded following placement of the tube on a magnetic rack, and the beads were washed with 1 mL of 80% ethanol in deionized water. Following careful removal of the wash solution, the beads were dried on the magnet for 2 minutes before being eluted with 24 *µ*L of deionized water. Subsequently, the absorbance profile and purity values of the purified RNA were assessed using a Nanodrop spectrophotometer. Finally, the sample was frozen at -80°C for storage until further downstream experiments, such as sequencing.

### 4.4 Nanopore direct RNA sequencing

#### RNA library preparation

All RNA libraries for direct RNA sequencing (DRS) were prepared according to the manufacturer’s protocol provided by Oxford Nanopore Technologies (ONT), with minor deviations for optimisation purposes. Initially, the RNA sample underwent sequential mixing with T4 DNA Ligase Buffer (Thermo Fisher Scientific), RT Adapter RTA, T4 DNA Ligase HC, and RNase In Plus for the ligation of RT Adaptor, followed by an incubation period as prescribed by ONT SQK002 protocol. Subsequently, reverse transcription was carried out by supplementing the mixture with 10 mM dNTPs, 5× SS UV First Strand Buffer, 0.1 M DTT, Nuclease-free water, and Superscript IV RT as per manufacturer’s instructions.

The resulting library underwent purification using Agencourt AMPure XP beads (Beckman), involving incubation, magnetic pelleting, ethanol washing, and resuspension steps. In the subsequent ligation step, the purified library was combined with RNA Adaptor RMX, T4 DNA Ligase HC, T4 DNA Ligase buffer, and molecular-grade water (mQ) to facilitate ligation with the library with a motor adaptor as per manufacturer’s instructions. A final cleanup procedure utilising Agencourt AMPure XP beads was performed, followed by elution in elution buffer ELB. All incubation durations, concentrations, and temperatures were rigorously maintained in accordance with SQK002 protocol provided by ONT. The resulting RNA library was subsequently directly used for loading or preserved at 4°C prior to loading.

#### Flow cell loading and live sequencing setup

The library loading procedure adhered to the protocol outlined by ONT for the MinION Mk1B or PromethION platform. The flow cell was inserted under the clip of the MinION, ensuring proper thermal and electrical contact. Quality control of the flow cell was performed, and the priming port was opened to remove any air bubbles. A priming mix, comprising Flush Tether and Flush Buffer (Activated Flush Buffer), was prepared, degassed, and loaded into the flow cell as per manufacturer’s instructions. Following this RNA library was prepared by mixing RNA Running Buffer, the ELB eluted RNA and molecular-grade water (mQ). Upon degassing, the library was gently loaded into the flow cell via the SpotON sample port in a dropwise manner. Finally, the priming port was closed, the SpotON sample port cover was replaced, and the MinION or PromethION was secured in place, following the recommended protocol guidelines. The DRS sequencing runs were conducted at room temperature (∼ 25°C) up to a maximum duration of 72 hours. Live basecalling was disabled during the run and the file format for data acquisition was set as FAST5 files for downstream preprocessing steps and subsequent bioinformatic analysis.

## 5 Conclusions

FracFixR provides a robust statistical framework for analyzing RNA-seq data from fractionation experiments, addressing the fundamental compositional challenge of recovering true fraction proportions from sequenced libraries. By modeling the relationship between whole and fractionated samples through non-negative linear regression, FracFixR enables accurate estimation of both global fraction weights and individual transcript distributions, including the often-overlooked “lost” fraction. The package’s ability to perform differential proportion testing between conditions makes it particularly valuable for identifying condition-specific changes in RNA localization or polysome association [15, 16]. FracFixR’s implementation as an open-source R package with a straightforward workflow ensures accessibility to the broader research community studying RNA compartmentalization, translation regulation, and subcellular localization. As RNA fractionation techniques continue to evolve and become more sophisticated, FracFixR provides an essential computational tool for extracting meaningful biological insights from these complex datasets while accounting for the inherent biases in library preparation and sequencing depth limitations.

## 6 Acknowledgments

The authors are grateful for the stimulating and insightful discussions around statistics and modeling held within the beautiful environment of Institut Montpelliérain Alexander Grothendieck (IMAG), University of Montpellier.

## 7 Data availability

The FracFixR R package is available from GitHub at https://github.com/Arnaroo/FracFixR. Synthetic tests data with full ground truth are available at the FracFixR GitHub page and can be flexibly generated using generate_data.py script included in the GitHub repository together with the generation instructions. The replicated DRS data for GM. REH and KOPN8 cell lines are available at NCBI SRA BioProject PRJNA1282458. The global fraction recovery synthetic mixes generated from these data are available as FASTQ and BAM files in the FracFixR GitHub site.

## References

[1] S. K. Archer, N. E. Shirokikh, T. H. Beilharz, and T. Preiss. Dynamics of ribosome scanning and recycling revealed by translation complex profiling. Nature, 535(7613): 570–574, 2016.

[2] S. Balzarini, R. Van Ende, A. Voet, and K. Geuten. A widely applicable and cost-effective method for specific rna–protein complex isolation. Scientific Reports, 13(1): 6898, 2023.

[3] Y. Benjamini and Y. Hochberg. Controlling the false discovery rate: a practical and powerful approach to multiple testing. Journal of the Royal statistical society: series B (Methodological), 57(1): 289–300, 1995.

[4] J. H. Bullard, E. Purdom, K. D. Hansen, and S. Dudoit. Evaluation of statistical methods for normalization and differential expression in mrna-seq experiments. BMC bioinformatics, 11: 1–13, 2010.

[5] A. R. Buxbaum, G. Haimovich, and R. H. Singer. In the right place at the right time: visualizing and understanding mrna localization. Nature reviews Molecular cell biology, 16(2): 95–109, 2015.

[6] H. Chassé, S. Boulben, V. Costache, P. Cormier, and J. Morales. Analysis of translation using polysome profiling. Nucleic acids research, 45(3):e15–e15, 2017.

[7] J. A. Christopher, A. Geladaki, C. S. Dawson, O. L. Vennard, and K. S. Lilley. Subcellular transcriptomics and proteomics: a comparative methods review. Molecular & Cellular Proteomics, 21(2): 100186, 2022.

[8] A. Conesa, P. Madrigal, S. Tarazona, D. Gomez-Cabrero, A. Cervera, A. McPherson, M. W. Szcześniak, D. J. Gaffney, L. L. Elo, X. Zhang, et al. A survey of best practices for rna-seq data analysis. Genome biology, 17: 1–19, 2016.

[9] M. D. Faye, T. E. Graber, and M. Holcik. Assessment of selective mrna translation in mammalian cells by polysome profiling. Journal of visualized experiments: JoVE, (92):52295, 2014.

[10] F. Hedayioglu, E. J. Mead, P. B. O’Connor, M. Skiotys, O. J. Sansom, G. R. Mallucci, A. E. Willis, P. V. Baranov, C. M. Smales, and T. von der Haar. Evaluating data integrity in ribosome footprinting datasets through modelled polysome profiles. Nucleic Acids Research, 50(19):e112–e112, 2022.

[11] C. E. Holt and S. L. Bullock. Subcellular mrna localization in animal cells and why it matters. Science, 326(5957): 1212–1216, 2009.

[12] A. Horvath, Y. Janapala, K. Woodward, S. Mahmud, A. Cleynen, E. E. Gardiner, R. D. Hannan, E. Eyras, T. Preiss, and N. E. Shirokikh. Comprehensive translational profiling and ste ai uncover rapid control of protein biosynthesis during cell stress. Nucleic Acids Research, 52(13): 7925–7946, 2024.

[13] N. T. Ingolia, G. A. Brar, S. Rouskin, A. M. McGeachy, and J. S. Weissman. The ribosome profiling strategy for monitoring translation in vivo by deep sequencing of ribosome-protected mrna fragments. Nature protocols, 7(8): 1534–1550, 2012.

[14] A. Jacobson and S. W. Peltz. Interrelationships of the pathways of mrna decay and translation in eukaryotic cells. Annual review of biochemistry, 65(1): 693–739, 1996.

[15] Y. Janapala, T. Preiss, and N. E. Shirokikh. Control of translation at the initiation phase during glucose starvation in yeast. International Journal of Molecular Sciences, 20(16): 4043, 2019.

[16] Y. Janapala, K. Woodward, J. Lee, M. Rug, T. Preiss, and N. E. Shirokikh. Rapid in vivo fixation and isolation of translational complexes from eukaryotic cells. Journal of Visualized Experiments, (178), 2021.

[17] L. F. Lareau, D. H. Hite, G. J. Hogan, and P. O. Brown. Distinct stages of the translation elongation cycle revealed by sequencing ribosome-protected mrna fragments. elife, 3:e01257, 2014.

[18] M. I. Love, W. Huber, and S. Anders. Moderated estimation of fold change and dispersion for rna-seq data with deseq2. Genome biology, 15: 1–21, 2014.

[19] J. S. Mattick. Non-coding rnas: the architects of eukaryotic complexity. EMBO reports, 2001.

[20] J. S. Mattick. Rna regulation: a new genetics? Nature Reviews Genetics, 5(4):316–323, 2004.

[21] J. S. Mattick, P. P. Amaral, P. Carninci, S. Carpenter, H. Y. Chang, L.-L. Chen, R. Chen, C. Dean, M. E. Dinger, K. A. Fitzgerald, et al. Long non-coding rnas: definitions, functions, challenges and recommendations. Nature reviews Molecular cell biology, 24(6): 430–447, 2023.

[22] A. E. Moor, M. Golan, E. E. Massasa, D. Lemze, T. Weizman, R. Shenhav, S. Baydatch, O. Mizrahi, R. Winkler, O. Golani, et al. Global mrna polarization regulates translation efficiency in the intestinal epithelium. Science, 357(6357): 1299–1303, 2017.

[23] M. D. Robinson, D. J. McCarthy, and G. K. Smyth. edger: a bioconductor package for differential expression analysis of digital gene expression data. bioinformatics, 26(1): 139–140, 2010.

[24] M. D. Robinson and A. Oshlack. A scaling normalization method for differential expression analysis of rna-seq data. Genome biology, 11: 1–9, 2010.

[25] N. E. Shirokikh, S. K. Archer, T. H. Beilharz, D. Powell, and T. Preiss. Translation complex profile sequencing to study the in vivo dynamics of mrna–ribosome interactions during translation initiation, elongation and termination. Nature protocols, 12(4): 697–731, 2017.

[26] E. C. Urdaneta, C. H. Vieira-Vieira, T. Hick, H.-H. Wessels, D. Figini, R. Moschall, J. Medenbach, U. Ohler, S. Granneman, M. Selbach, et al. Purification of crosslinked rna-protein complexes by phenol-toluol extraction. Nature communications, 10(1): 990, 2019.

[27] E. Villanueva, T. Smith, M. Pizzinga, M. Elzek, R. M. Queiroz, R. F. Harvey, L. M. Breckels, O. M. Crook, M. Monti, V. Dezi, et al. System-wide analysis of rna and protein subcellular localization dynamics. Nature Methods, 21(1): 60–71, 2024.

[28] J. Wang, M. Horlacher, L. Cheng, and O. Winther. Rna trafficking and subcellular localization—a review of mechanisms, experimental and predictive methodologies. Briefings in bioinformatics, 24(5):bbad249, 2023.

[29] Z. Wang, M. Gerstein, and M. Snyder. Rna-seq: a revolutionary tool for transcriptomics. Nature reviews genetics, 10(1): 57–63, 2009.

